# Disentangling object category representations driven by dynamic and static visual input

**DOI:** 10.1101/2022.05.03.490462

**Authors:** Sophia Robert, Leslie G. Ungerleider, Maryam Vaziri-Pashkam

**Affiliations:** Lab of Brain and Cognition, National Institute of Mental Health, Bethesda, MD, USA; Department of Psychology and Neuroscience Institute, Carnegie Mellon University, Pittsburgh, PA, USA

## Abstract

Humans can label and categorize objects in a visual scene with high accuracy and speed—a capacity well-characterized with neuroimaging studies using static images. However, motion is another cue that could be used by the visual system to classify objects. To determine how motion-defined object category information is processed in the brain, we created a novel stimulus set to isolate motion-defined signals from other sources of information. We extracted movement information from videos of 6 object categories and applied the motion to random dot patterns. Using these stimuli, we investigated whether fMRI responses elicited by motion cues could be decoded at the object category level in functionally defined regions of occipitotemporal and parietal cortex. Participants performed a one-back repetition detection task as they viewed motion-defined stimuli or static images from the original videos. Linear classifiers could decode object category for both stimulus formats in all higher order regions of interest. More posterior occipitotemporal and ventral regions showed higher accuracy in the static condition and more anterior occipitotemporal and dorsal regions showed higher accuracy in the dynamic condition. Significantly above chance classification accuracies were also observed in all regions when training and testing the SVM classifier across stimulus formats. These results demonstrate that motion-defined cues can elicit widespread robust category responses on par with those elicited by luminance cues in regions of object-selective visual cortex. The informational content of these responses overlapped with, but also demonstrated interesting distinctions from, those elicited by static cues.

**Significance Statement:** Much research on visual object recognition has focused on recognizing objects in static images. However, motion cues are a rich source of information that humans might also use to categorize objects. Here, we present the first study to compare neural representations of several animate and inanimate objects when category information is presented in two formats: static cues or isolated dynamic cues. Our study shows that while higher order brain regions differentially process object categories depending on format, they also contain robust, abstract category representations that generalize across format. These results expand our previous understanding of motion-derived animate and inanimate object category processing and provide useful tools for future research on object category processing driven by multiple sources of visual information.

## Introduction

Humans can categorize objects with striking speed and accuracy. Previous research on the neural basis of visual object recognition has largely focused on the processing of static features from images along the ventral visual hierarchy of the primate brain (reviewed in Peissig & Tarr, 2007). However, real-world scenes are not static. In fact, decades of behavioral research have shown that motion cues can contain category-relevant information that humans use to make judgements about objects. Behavioral studies using point-light displays (PLDs, Johansson, 1973; Johansson, 1976) have established that, even with the impoverished motion information available in PLDs, humans can quickly perceive a moving person, identify the action being performed, and even determine the actor’s age, gender, and affect (e.g., Barclay et al., 1978; Bassili, 1978; Cutting and Kozlowski, 1977; Dittrich et al., 1996).

The majority of biological motion research has focused on the perception of human motion due to the significant role that it plays in our social lives. However, our sensitivity to information in motion cues is not restricted to perceiving humans. Humans can also infer animacy and complex social relations from the movements of basic geometric shapes (Schultz & Bülthoff, 2013; Heider & Simmel, 1944; Scholl & Gao, 2013) and can recognize animal categories such as chickens, dogs, horses and cats in PLDs (Mitkin & Pavlova, 1990; Mather & West, 1993; Pinto & Shiffrar, 2009; Pinto, 1994; Pavlova et al., 2001).

Investigations of the neural underpinnings of object categorization from motion information with neuroimaging have identified the superior temporal sulcus (STS) as a key region involved in processing biological motion. The STS has been shown to track animacy signals in motion cues from simple shapes and to process dynamic movements of human faces and bodies (Schultz & Bulthoff, 2013; Hirai & Hiraki, 2006; Pitcher et al. 2011, Pavlova et al., 2004). Neuropsychological studies have also suggested the involvement of parietal regions in the integration of motion and form information during form-from-motion identification tasks (Schenk & Zihl, 1997).

Despite extensive research into neural substrates of human motion processing (Giese, 2013), there have been comparatively few studies that have investigated how non-human motion is processed in the brain. Previous studies suggest preferential processing of human motion over that of one or two other classes, e.g., mammals or tools, in regions in lateral occipito-temporal cortex (LOTC) including the posterior STS (Papeo et al., 2017), human middle temporal complex (Kaiser et al., 2012), and fusiform gyrus (Grossman & Blake, 2002), as well as the inferior parietal lobe, inferior frontal gyrus (Saygin et al., 2004), the posterior and anterior cingulate cortices and the amygdala (Bonda et al., 1996; Ptito et al., 2003).

The limited neuroimaging studies that have directly compared object representations driven by motion to those driven by static images have focused on human (or monkey) faces and bodies (Furl et al., 2012; Hafri et al., 2017; Pitcher et al., 2011) or have only compared humans with tools (Beauchamp et al., 2003). Furthermore, these studies (with the exception of Beauchamp et al., 2003), have used videos containing both static and dynamic cues as their dynamic condition and thus have not been able to carefully separate the contributions of motion- and image-information to the responses. Thus, a systematic comparison of several object category representations driven by isolated motion and static cues has yet to be undertaken.

Here, we devised a novel method to generate stimuli that only contained motion cues. We extracted motion signals from videos of objects and simulated object movements using flow fields of moving dots. We first demonstrated that humans can recognize a wide variety of animate and inanimate objects in our dynamic stimuli. We then used these stimuli, along with static images, in an fMRI study to compare object category representations derived from dynamic and static cues in occipito-temporal and parietal regions of interest across visual cortex.

## Materials and Methods

### Stimuli

#### Stimulus creation pipeline

Eight categories were selected to sample a wide range of animate and inanimate object categories: human, non-human mammal, bird, reptile, vehicle, tool, pendulum/swing, and ball. We sought videos of objects performing a wide range of movements. Video clips were downloaded from various sources on the Internet or shot with in-house equipment in accordance with the following criteria: 1) contained a single moving object, 2) contained the entire object in frame without occlusion, 3) shot without camera movement (no zooming, panning, tracking), 4) contained no movement in the background, and 5) lasted at least 3 seconds.

We used in-house Matlab code, the Psychtoolbox extension, and in-house python code to generate moving dot patterns that followed the movement of the objects in the videos. To do this, first, all videos were trimmed to 3 seconds, cropped with a 3:2 x/y aspect ratio to center the object, and resized to 720 × 360 pixel resolution. Videos with 30 frames per second were then up-sampled so that all videos had a frame rate of 60 fps. The local, frame-by-frame motion of the objects in each video in x and y directions was then extracted using the Farneback optical flow algorithm (Farneback, 2003).

Next, object movements extracted from the full videos were projected on moving dot patterns. To create the moving dot stimuli, 2500 white dots (2 pixel diameter) were randomly initialized on a grey background (360 × 720 pixels). Dots that fell within pixels with nonzero motion vector values were moved in the direction and magnitude specified by the extracted motion matrix in the next frame. The lifetime (number of contiguous frames of movement) of any dot was randomly sampled from a uniform distribution between 1 and 17 frames. The lifetime value decreased on every frame. If the lifetime of a dot reached 0 or they reached the boundaries of the frame, they were reinitialized with a lifetime of 17 frames.

The number of dots for a given frame and their lifetime was set to mitigate the formation of dot clusters that could induce perception of an edge in individual frames of the video. The frames were qualitatively examined to see if they induced a perception of any kind of edge or form. Videos that produced such artifacts were removed from the stimulus set. For the fMRI experiment, these moving dot videos were rendered live for each trial so that the dot initializations were always random.

#### Stimulus Validation Experiment

To ensure that the stimuli contained clear category information, we conducted an online experiment. 430 participants (223 women, aged 18-65) were recruited on Amazon Mechanical Turk to perform an object categorization task on the dynamic stimuli. Participants each performed between 10-11 trials. For each trial, participants were asked 3 questions about the object in a looped video: 1) whether the object in the video was of an animal or non-animal, 2) which of 8 listed categories the object belonged to, and 3) whether they could label the object. If subjects responded ‘yes’ for the third question, they were required to type the label in a response text box. Each of the three questions contained an “I don’t know” option. Subjects had to answer all three questions to complete each trial.

Overall, subjects categorized objects based on their motion in the moving dot stimuli with an average accuracy of 76% (202 total videos). The three animate (human, mammal, reptile) and three inanimate (tool, ball, pendulum/swing) categories with the highest accuracy were used for the fMRI experiment. For each category, the 6 videos with the highest accuracy were selected (mean accuracy = 96%).

The overall ‘motion energy’ of each video was calculated by averaging the motion vectors across all pixels in all frames. Non-zero motion vectors were also used to calculate the average non-zero ‘motion energy’. The average overall and non-zero motion energy for the 6 videos in each category were entered into pairwise two-sample heteroscedastic t-test comparisons to ensure that there were no significant differences between categories for either metric. Neither the overall nor the non-zero motion energies were significantly different across categories (all *p*s > 0.05, even without correction for multiple comparisons).

After the dynamic video stimulus set was finalized, the static image stimulus set was generated by randomly selecting three frames of the full form video from which the moving dot stimulus was created. The frame with the object in clearest view was selected and further processed to extract the object from the frame. For the fMRI experiment, the isolated object was pasted onto a background of 2500 randomly initialized white dots on a grey background, to mimic a frame of the dynamic moving dot stimuli.

### Functional MRI experiment

#### Participants

Fifteen healthy human subjects (six women, age range 19-42) with normal or corrected to normal vision were recruited for the fMRI experiment. Participants were brought in for a 2 h fMRI session that included the main experiment and three localizer tasks. Prior to entering the scanner, all participants practiced the tasks for the main experiment and localizer runs and underwent a short behavioral task to familiarize themselves with the stimuli. All subjects provided informed consent and received compensation for their participation. The experiments were approved by the NIH ethics committee.

#### Training Session

The independent norming study performed with mTurk demonstrated that people can recognize the objects in these stimuli with high accuracy after minimal instruction. However, to avoid introducing any random factors across subjects and differential processing during the first run of the session relative to the rest, participants participated in a training session prior to entering the scanner. During the training session, they familiarized themselves with the 36 dynamic stimuli and were subsequently tested to ensure accurate recognition. Each video was shown on loop until subjects could verbally report which of the 6 categories the object belonged to. If the subject categorized the object correctly, the experimenter advanced to the next stimulus; incorrect categorizations were verbally corrected by the experimenter. After all stimuli had been verbally categorized, subjects underwent a testing session. In each trial, a random video was shown once without looping, followed by a grey screen with 6 category labels placed in a circle around the center of the screen. Subjects were instructed to categorize the object in the video by clicking on the corresponding category label. No feedback was provided during the testing session. If a subject performed above 90% accuracy, they continued on to the fMRI experiment. The training and testing session took no longer than 15 minutes. Subjects required little to no correction during the training session and performed with an average of 99% accuracy in the test session on the first iteration (n = 13, data for two subjects were lost due to technical problems).

#### MRI Methods

MRI data were collected from a Siemens MAGNETOM Prisma scanner at 3 Tesla equipped with a 32-channel head coil. Subjects viewed the display on a BOLDscreen 32 LCD (Cambridge Research Systems, 60 Hz refresh rate, 1600 × 900 resolution, at an estimated distance of 187 cm) through a mirror mounted on the head coil. The stimuli were presented using a Dell laptop with MATLAB and Psychtoolbox extensions (Brainard, 1997; Kleiner, Brainard, & Pelli, 2007).

For each participant, a high resolution (1.0 × 1.0 × 1.0 mm) T1-weighted anatomical scan was obtained for surface reconstruction. All functional scans were collected with a T2*-weighted single-shot, multiple gradient-echo EPI sequence (Kundu et al., 2012) with a multiband acceleration factor of 2 slices/pulse. 50 slices (3 mm thick, 3 × 3 mm^2^ in-plane resolution) were collected to cover the whole brain (TR 2 s, TE = 12 ms, 28.28 ms, 44.56 ms, flip angle = 70°, FoV = 216 mm).

#### Experimental Design

##### Main Experiment

The main task of the experiment included 6 categories: human, mammal, reptile, tool, pendulum/swing, and ball and 2 stimulus conditions: dynamic (moving dot videos) and static (object images pasted on dot background). Both dynamic and static stimuli were presented at the same size and location (subtending 9.6° x 4.8° visual angle). We used a block design to present alternating blocks of dynamic and static stimuli while also alternating between animate and inanimate blocks. The order of the six categories and the two formats were counterbalanced within and across runs. Four different counterbalancing designs were created and each subject was randomly assigned one of the designs.

Each run contained 12 condition blocks, one for each condition (2 formats x 6 categories), began with an initial fixation block of 8 s, and ended with a final fixation of 12 s. Each condition block began with an 8 s fixation period in which a red fixation dot (5 pixels in radius) was shown on a grey background. The fixation period was then followed by the stimulus presentation period in which 4 stimuli were presented from the same condition, each for 2.8 s followed by a 200 ms inter-stimulus interval, resulting in 12 s of stimulus presentation. The duration of each condition block was 20 s (8 s fixation and 12 s stimulus presentation). For each run, the 12 condition blocks and the initial and final fixation blocks lasted 252 s (4 min 12 s). Each participant completed 12 runs.

To maintain their attention, subjects were given a one-back repetition detection task in which they were instructed to press a button on an MRI-compatible button box (fORP, Cambridge Research Systems) to indicate detection of a repeated stimulus within each block. There was one stimulus repetition per block and the repeated stimulus of each block type was changed across runs. Because there were only 3 unique trials per block but each condition had 6 unique stimuli, half of the stimuli of each category were shown on odd runs and the other half were shown on the even runs. These blocks were later combined during analysis. Average performance on this task was 94%. To ensure proper fixation, eye movements were monitored using an ASL eye-tracker.

##### Object Localizer task

To localize functional ROIs in ventral and lateral occipito-temporal cortex, we presented images of objects in 6 conditions: faces, scenes, head-cropped bodies, central objects, peripheral objects (4 objects per image), and phase-scrambled objects in a block design paradigm. Subjects were instructed to fixate while 20 images were presented in each block for 750ms with a 50ms fixation screen in between. Each block lasted 16 s and was repeated 4 times per condition. Each run started with a 12s fixation period. Additional 8 s fixation periods were presented after every 5 blocks. Total run duration was 436 s (7 min 16 s). Subjects performed a motion detection task. During each block, a random image would jitter by rapidly shifting 4 pixels back and forth horizontally from the center of the screen. Subjects indicated detection of motion with a button press. Each participant completed 1-2 runs of this task.

##### Motion localizer task

To localize functional ROIs related to the perception of biological and non-biological motion, we presented blocks of point light display (PLD) videos of humans performing various actions in four conditions: 1) biological motion: normal PLD video (e.g. walking, riding a bicycle), 2) random motion: the points in the PLD were spatially scrambled in each frame, 3) translation: randomly positioned dots translated across the screen in a random direction with the speed set to the average speed of the movement from the PLD videos, and 4) static: a random frozen frame of the PLD was shown as an image. There were 8 exemplars per condition, each presented for 1.5 s followed by a 500 ms interstimulus fixation period. Each block lasted 16 s and was presented 4 times per condition. Each run began with a 6s fixation period and 8 s fixation periods were interspersed between each block making the total run duration 422.7 s (7 min 3 s). Subjects performed a one-back repetition detection task, in which they indicated detection of a repeated stimulus during each block by pressing a button. Each subject completed 1-2 runs of this task.

##### Topographic mapping

Topographic visual region V1 was mapped using 16 s blocks of a vertical or horizontal polar angle wedge with an arc of 60° flashing black and white checkerboards at 6 Hz. During the stimulus blocks, subjects fixated on a red fixation dot (5 pixel radius) and detected a dimming on the wedge, that occurred randomly either at the inner, middle, or outer ring of the wedge at 4 random times within the 16 s block. There was a 16 s fixation period after each block and each run began with a 16 s period of fixation. Each run lasted 272 s (4 min and 40 s), and subjects completed 1-2 runs of this task.

#### Data Analysis

fMRI data were analyzed using AFNI (Cox, 1996) and in-house MATLAB codes. The data were pre-processed by removing the first 2 TRs of each run, motion correction, slice timing correction, smoothing with 5mm FWHM, and intensity normalization. The EPI scans were registered to the anatomical volume. The three echoes were combined using a weighted average (Posse et al., 1999; Kundu et al., 2012). TRs with motion exceeding 0.3 mm as well as outliers were excluded from further analysis. A general linear model analysis with 12 factors (2 stimulus conditions x 6 categories) was used to extract t-values for each condition in each voxel. The 6 degrees of freedom movement parameters was used as an external regressor. To account for the effect of residual autocorrelation on statistical estimates, we applied a generalized least squares time series fit with restricted maximum likelihood (REML) estimation of the temporal auto-correlation structure in each voxel. The t-values were calculated across all runs for the univariate analysis and per-run for the multivariate analysis.

#### ROI Definition: Group-constrained subject specific method

We used a systematic, unbiased method for creating individualized regions of interest constrained by group responses to our localizer experiments, basing our approach on a method of region of interest definition developed by Kanwisher and Fedorenko (described in Pitcher et al., 2011).

First, t-values were extracted from generalized linear models (GLMs) of individual activation maps from the localizer experiments. All subjects’ statistical activation maps (N = 15) were converted to Talairach space. For each subject, the individual localizer contrast maps were thresholded at *p* < 0.0001. Group overlap proportion maps were then created for each contrast.

Second, we thresholded the group proportion maps for each contrast separately to counteract contrast- or localizer-specific differences in spatial variability or overall activation. The thresholds for specific contrast maps were as follows: For the object localizer experiment, the thresholds were N ≥ 0.7 for objects vs scrambled (lateral occipital, LO; posterior fusiform sulcus, pFS), N ≥ 0.5 for bodies vs objects (extrastriate body area, EBA), and N ≥ 0.25 for peripheral objects vs scrambled (inferior intraparietal sulcus, infIPS). For the biological motion experiment, the threshold for biological motion vs translation was N ≥ 0.5 (lateral occipito-temporal biomotion region, LOT-biomotion). For the retinotopy experiment, positive and negative maps were created separately and thresholded at N ≥ 0.5.

Third, we used a Gaussian blur of 1mm FWHM. The blurred maps were then clustered using the nearest neighbors method and a minimum cluster size of 20 voxels. For V1, positive and negative maps were clustered separately and then combined with a step function. Two steps were required to finalize the group-constrained ROIs. Anatomical landmarks were used to separate pFS from LO, and LO from infIPS. V1 was separated from V2 using a hand-drawn region based on the group map. All ROIs were then selected to have no overlapping voxels.

The final nonoverlapping group-constrained ROIs were made subject specific by creating masks based on the individual subject’s activity during the localizer experiments (localizer contrast threshold: p < 0.05). For example, for each subject’s EBA, the group-constrained EBA was masked by the subject’s response to bodies > objects with a threshold of p < 0.05. If this process did not yield an ROI with at least 100 voxels across the two hemispheres, the ROI was instead created with a mask made from the mean response during the main experiment (task vs fix, p < 0.0001 uncorrected).

The supramarginal (SMG) region of interest was anatomically defined using a Freesurfer parcellation (Desikan et al, 2006). To make the subject specific supramarginal ROIs, individual masks were made from the mean response during the main experiment (task vs fixation, p < 0.0001 uncorrected) and intersected with the template SMG region.

#### Univariate analysis

To calculate the average fMRI response per condition for each ROI, using a general linear model analysis, whole brain t-value maps were extracted for each of the 12 conditions and masked with a task > fixation threshold of p < 0.0001 for each subject. The group-constrained subject-specific ROIs were intersected with these maps, resulting in a t-value response per voxel in each ROI for all 12 conditions in each subject. The average responses for four conditions were then calculated from these ROI responses: dynamic animate, dynamic inanimate, static animate, and static inanimate. The animacy preference in each ROI was calculated as the difference between the animate and inanimate conditions, separately for the static and dynamic stimulus formats. One-sample and paired t-tests were conducted to determine respectively: 1) if the animacy preference in each ROI and each format was significantly different from 0, and 2) if the animacy preference was significantly different across stimulus formats within each ROI. All t-tests were corrected for multiple comparisons with False Discovery Rate correction (Benjamini and Hochberg, 1995) across ROIs.

#### Multivariate pattern analysis (MVPA)

We performed multivariate pattern analyses to investigate whether object category information was present in the fMRI responses to the dynamic and static stimuli. We extracted t-values in each voxel for every condition in each run using a GLM analysis. To perform pairwise object category decoding, we used a linear support vector machine classifier (SVM; Chang and Lin, 2011) with feature selection. The SVM was trained using leave-one-out cross validation on data that was normalized with z-scoring to avoid magnitude differences between conditions. Using t-tests, we calculated the top 100 most informative voxels per ROI (Mitchell et al., 2004) to equate the number of voxels analyzed per ROI and facilitate comparisons between them. This feature selection was performed separately for each iteration of training. Results did not qualitatively change when the analysis was performed without feature selection.

We trained and tested the linear SVM in two conditions: 1) within-classification, in which the SVM was trained and tested on the same stimulus format, and 2) cross-classification, in which SVM was trained in one stimulus format and tested on the other format. The classification was performed on all unique pairs of object categories to obtain classification accuracy matrices. The off-diagonal values of the matrices were averaged to produce two within-format and two cross-format average object category decoding accuracies per subject. The two cross-format values were then averaged to obtain one cross-classification accuracy. One-sample and paired t-tests were conducted to determine respectively: 1) if the decoding accuracy in each ROI and each format was significantly different from chance (0.5), and 2) if the decoding accuracy was significantly different across stimulus formats within each ROI. All p-values listed from t-tests and ANOVAs were corrected for multiple comparisons with False Discovery Rate correction across ROIs (Benjamini and Hochberg, 1995). For ANOVAs, effect sizes were calculated with generalized eta squared 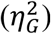, for the one sample and paired t-tests, Cohen’s *d* was used.

#### Multidimensional scaling of fMRI responses

To visualize how stimulus format and object category impact the responses in our regions of interest, we quantified the similarities between the patterns of fMRI responses to the 12 conditions in each ROI by calculating all pairwise Euclidean distances. The individual subject Euclidean distances per ROI were averaged across subjects to create group Euclidean distances, which will be referred to as the fMRI-Euclidean matrix. We then visualized these similarities by applying classical multidimensional scaling (Shepard, 1980) on the fMRI-Euclidean matrix and plotting the first two dimensions for each ROI.

We measured the reliability of the fMRI-Euclidean matrix by performing a permutation analysis wherein the individual subject matrices were split into two groups, averaged to create two group matrices, and then correlated to get a measure of the split-half reliability. Correlations for every possible combination of subjects in the two groups were measured and averaged to produce a final reliability score. The reliabilities of the dynamic and static fMRI-Euclidean matrices were evaluated separately.

### Object similarity behavioral experiment

353 participants (32% female among the 85% who responded to the demographic survey) were recruited on Amazon Mechanical Turk to perform an object similarity task on the dynamic or static stimuli. All participants were located in the United States.

For each trial, participants were presented with three stimuli on a grey screen and were instructed to select the ‘odd-one-out’ stimulus (the stimulus that was most distinct among the three) by clicking on it. Dynamic and static stimuli were tested separately. Participants performed blocks of 15 trials to complete the task and were permitted to perform more than one block. To ensure data quality, trials with RTs smaller than 0.6 s and 1.2 s and larger than 10 s or 20 s were removed for the image and video tasks, respectively. These cutoffs were decided based on the distributions of RTs. If 5 or more trials in a block were eliminated, the entire block (or HIT in mTurk terminology) was removed. The eliminated blocks were resubmitted to mTurk to ensure that we had at least 2 repetitions for each unique triplet allowing for 68 trials for each pair of stimuli.

To build a dissimilarity matrix based on the odd-one-out image and video tasks, a response matrix of the pairwise dissimilarity judgments was constructed for each task by treating each triplet as three object pairs and assigning 1’s to dissimilar pairs (i.e. the two pairs that included the selected odd object) and a 0 to the similar pair (i.e. the pair that did not include the selected odd object). We also constructed a count matrix to determine how many times each pair was shown together in a triplet. By dividing the response matrix by the count matrix, we obtained a dissimilarity matrix with values ranging from 0-1 with higher values denoting higher dissimilarity. To produce a category level behavioral dissimilarity matrix, we took the off-diagonal upper triangle of the 36 × 36 matrix and averaged the item distances that belonged to the same category, resulting in a 6 × 6 matrix, which will be referred to as the behavioral-dissimilarity matrix. The diagonal was nonzero due to nonzero distances between exemplars within each category. Only the off-diagonal of this matrix was used in further analyses.

To gauge the stability of the behavioral-dissimilarity matrix, we performed a split-half reliability analysis. Because each subject only saw a small set of all possible triplets, instead of splitting the data by subject, we split based on repeats of stimulus pairs (3 pairs per triplet) into two groups. The binary similarity values for all pairs were correlated across the two groups to produce a measure of reliability of the similarity judgments.

#### Multi-dimensional scaling and hierarchical clustering of object similarity responses

We visualized the structure of the object similarity judgments from the odd-one-out tasks at the category level using classical multidimensional scaling on the behavioral-dissimilarity matrices of the dynamic and static stimuli separately (Shepard, 1980). The two behavioral-dissimilarity matrices were also correlated to quantify their degree of similarity. To investigate the structure of the object similarity judgments at the exemplar level, we used a hierarchical or agglomerative clustering algorithm available in the Python package *scipy* (Virtanen et al., 2020) on the dynamic and static behavioral-dissimilarity matrices separately. For visualization purposes, images of the individual exemplars, which were adapted from the static stimuli used in the experiment, were included under the resultant dendrograms for both static and dynamic conditions (note that dynamic stimuli are not recognizable in static frames).

#### Brain-behavior correlation

To determine the relationship between the multivariate information for the six categories in each region of interest (fMRI-Euclidean matrix) with behavioral assessments of the category similarity (behavioral-dissimilarity matrix), we correlated the two measures. For each subject, the off-diagonal of the fMRI-Euclidean matrix was correlated with the off-diagonal behavioral-dissimilarity matrix using Pearson’s linear correlation coefficient, separately for the dynamic and static experiments. The correlations were then averaged across subjects. The noise ceiling of these correlations was then calculated for each ROI as the square root of the product of the reliabilities of the fMRI-Euclidean matrix and the behavioral-dissimilarity matrix. As the reliability of the behavioral-dissimilarity matrix was calculated with only one split, the standard error of the noise ceiling was calculated based on the mean and standard deviation of the reliability scores generated on each permutation of the fMRI-Euclidean reliability analysis.

#### Brain-optic flow correlation

To ensure that optic flow information from the six object categories was not predictive of the multivariate fMRI responses in any of the regions of interest, we performed a control analysis. We first calculated the Euclidean distances between the dynamic stimulus information of each category by vectorizing the 4-dimensional stimuli (x-coordinates, y-coordinates, x- and y-magnitudes of optic flow, and time) and averaging the distances between stimuli of the same category, creating the optic flow-Euclidean matrix. We then correlated the optic flow-Euclidean matrix with the dynamic fMRI-Euclidean matrix of each ROI for each subject. The correlations were averaged across subjects to generate group mean correlations and one-sampled t-tests were used to determine whether any positive correlations were significantly above zero.

## Results

### Effect of stimulus format on univariate animacy preference

We first looked at the mean amplitude of responses to the two superordinate object categories (animate/inanimate) in the two stimulus formats (static/dynamic). We extracted individual subjects’ t-values from the GLM analysis and averaged the response for the three animate and the three inanimate categories within each image format to get 4 values per subject. Figure 2 shows the pooled results of this analysis across subjects. A two-way ANOVA with stimulus format and animacy as factors showed a significant main effect of stimulus format in all ROIs (*f*s > 7.26, *p*s ≤ 0.02 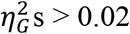) with higher response amplitude in the dynamic compared to the static condition. A main effect of animacy was also found in LO, pFS, EBA, LOT-biomotion, and left SMG (*f*s > 7.68, *p*s < 0.03, 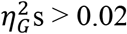), but not in V1, infIPS, or right SMG (*f*s < 3.38, *p*s > 0.12, 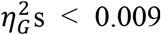). For the four ventrotemporal cortical areas, average responses were significantly higher for the animate object categories, while in left SMG the average response was higher for the inanimate object categories. The pattern of responses in SMG was not solely driven by the tool category as removing tools from the inanimate objects did not qualitatively change the results (data not shown).

**Figure 1.**
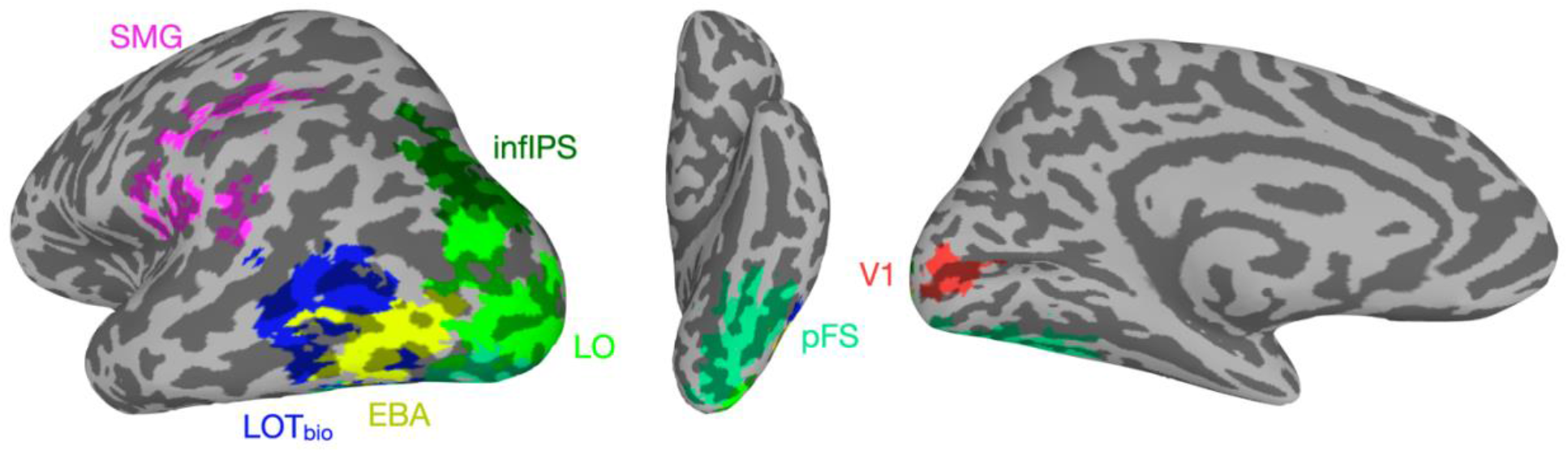
Regions of interest of a single example subject generated by the group-constrained single-subject method. The supramarginal area (SMG) is colored in pink, the inferior intraparietal sulcus (infIPS) is colored in dark green, the lateral occipital complex (LO) is colored in light green, the extrastriate body area (EBA) is colored in yellow, the biological motion related lateral occipito-temporal area (LOT-bio) is colored in dark blue, the posterior fusiform sulcus (pFS) is colored in teal, and primary visual cortex (V1) is colored in red.

**Figure 2.**
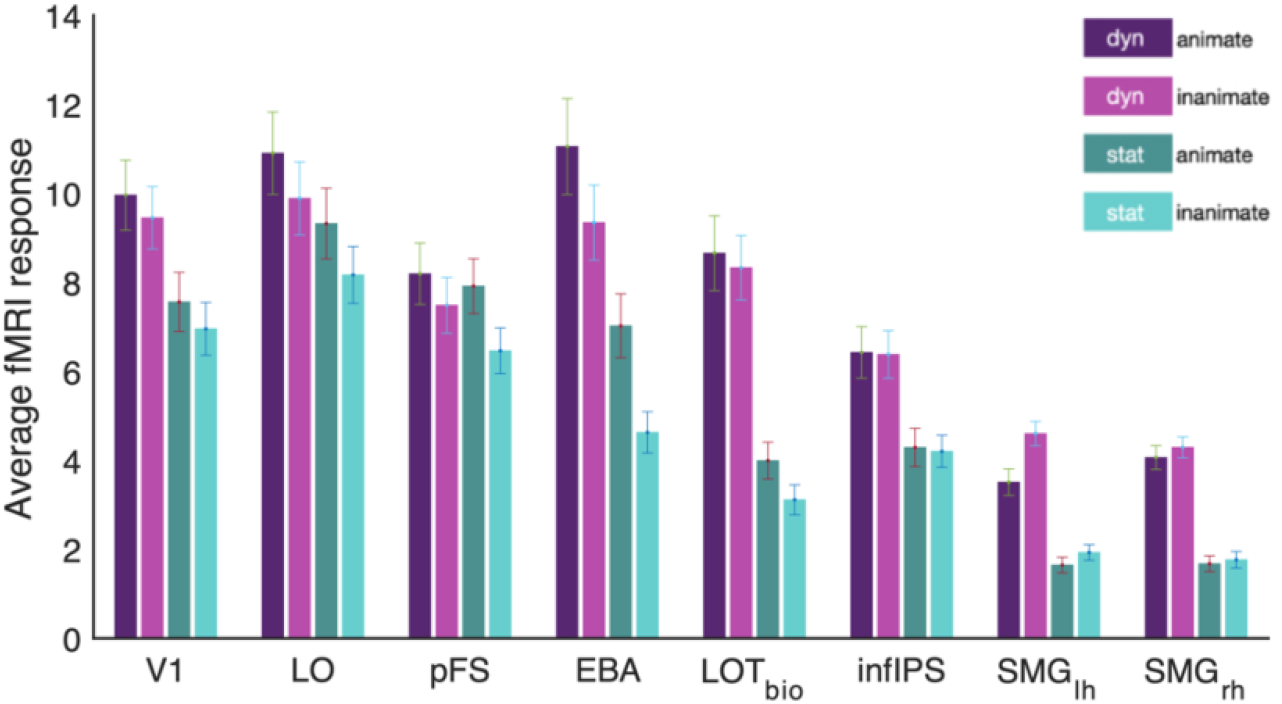
Univariate fMRI responses to dynamic and static stimuli averaged within animate and inanimate categories for each region of interest. Results do not qualitatively differ when removing the human and tool categories from the analysis. Error bars represent standard errors.

To better visualize and investigate the interaction between stimulus format and animacy, we subtracted inanimate responses from animate responses to produce a measure of animacy preference within each stimulus format (Figure 3). Unpaired t-tests evaluating animacy preference against 0 revealed that there was no animacy preference in V1, inferior IPS, and the right SMG area in either stimulus format (dynamic: *t*s < 1.56, *p*s > 0.21, Cohen’s *d*s < 0.42, static: *t*s < 0.76, *p*s > 0.55, Cohen’s *d*s < 0.20). In contrast, for both stimulus formats, LO, pFS, and EBA showed a preference for animate categories (dynamic: *t*s > 3.15, *p*s < 0.02, Cohen’s *d*s > 0.84, static: *t*s > 5.05, *p*s < 0.0002, Cohen’s *d*s > 1.35) while left SMG preferred inanimate categories (dynamic: *t*(14) = 5.59, *p* = 0.0005, Cohen’s *d* = 1.49). LOT-biomotion had significant preference for animate categories in the static (*t*(14) = 3.97, *p* = 0.003, Cohen’s *d* = 1.06) but not in the dynamic condition (*t*(14) = 1.14, *p* = 0.31, Cohen’s *d* = 0.31). All regions showed a preference in the same direction for dynamic and static conditions.

**Figure 3.**
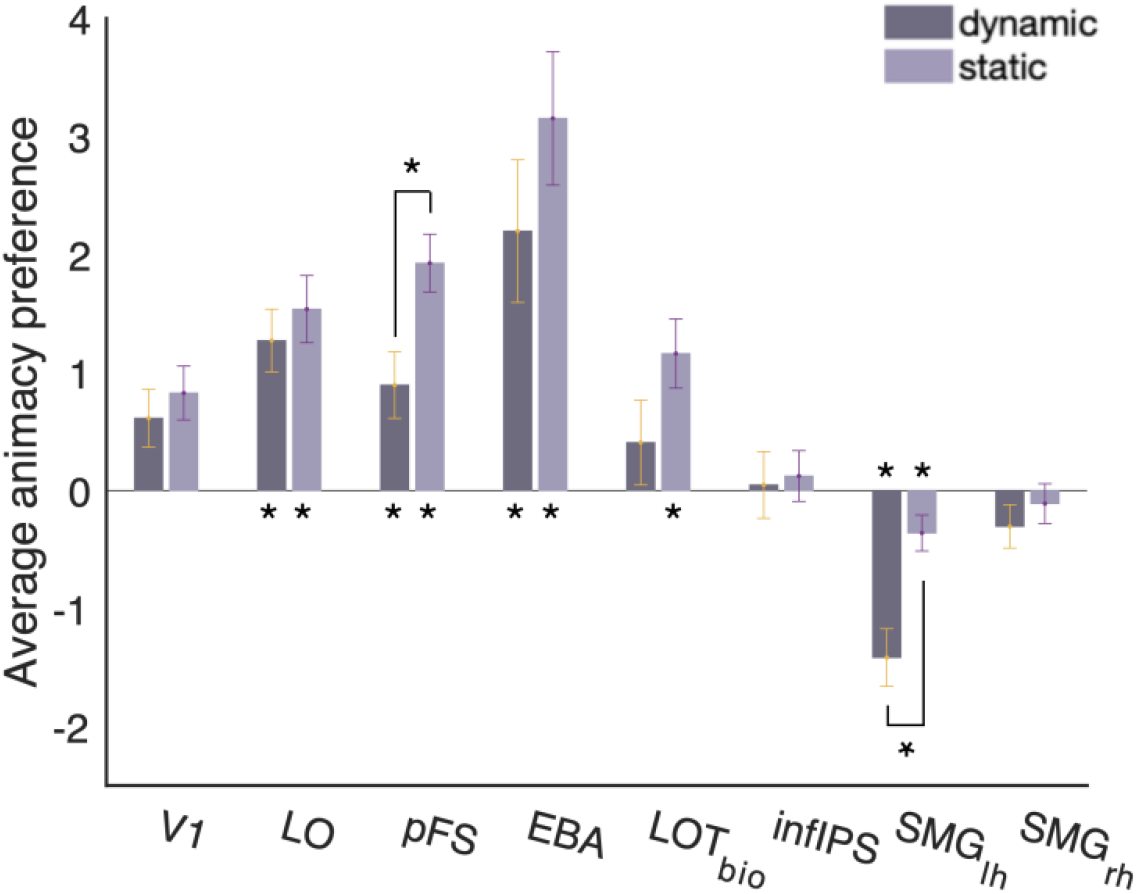
Univariate fMRI response preference for animate compared to inanimate object categories in dynamic and static stimuli for each region of interest. \**p*s < 0.05. Error bars represent standard errors.

pFS and left SMG further showed a significant difference in the magnitude of their animacy preference across formats. pFS, a ventral region known to be involved in object recognition, showed a stronger preference for animate object stimuli in the static compared to the dynamic condition (paired t-test: *t*(14) = 3.07, *p* = 0.03, Cohen’s *d* = 0.79), while left SMG, a parietal region thought to be involved in tool processing and action observation had a stronger preference for inanimate object stimuli in the dynamic compared to the static condition (paired t-test: *t*(14) = 3.73, *p* = 0.02, Cohen’s *d* = 0.96). These significant interactions between stimulus format and animacy preference suggest that the category preference responses in pFS and left SMG are modulated by the format through which the category information is provided. The most ventral region, pFS, is more sensitive to static form presentations of animate objects and the most dorsal lateral region, left SMG, is more sensitive to dynamic motion information about inanimate objects.

### Effect of stimulus format on multivariate object category representations

We next examined the multivariate patterns of each of our regions of interest to further explore how object category information is represented in the brain when sourced from dynamic movements and static images. We first sought to test if each of our regions contained information about the 6 object categories within each stimulus format. To do this, we calculated average pairwise classification accuracy for the 6 object categories for the static and dynamic conditions using a linear SVM classifier (Chang and Lin, 2011). Figure 4a shows the pooled results of this analysis across subjects. Unpaired t-tests revealed that the object categories were decoded significantly above chance in both dynamic and static formats in all regions but V1 (dynamic: *t*s > 7.04, *p*s < 0.00001, Cohen’s *d*s > 1.82; static: *t*s > 2.73, *p*s < 0.02, Cohen’s *d*s > 0.71). In V1, significant decoding was only found in the static stimulus condition (static: *t*(14) = 8.31, *p* = 0.00001, Cohen’s *d* = 2.15; dynamic: *t*(14) = 2.05, *p* = 0.06, Cohen’s *d* = 0.53). In all regions but infIPS, there were significant differences between the decoding accuracies across stimulus format (infIPS: *t*(14) = 0.59, *p* = 0.57, Cohen’s *d* = 0.15). In V1, LO, pFS, and EBA decoding accuracies were higher in the static condition than the dynamic (*t*s > 2.32, *p*s < 0.001, Cohen’s *d*s > 0.60), while in LOT-biomotion and bilateral SMG, decoding accuracies were higher in the dynamic condition (*t*s > 3.24, *p*s < 0.008, Cohen’s *d*s > 0.84).

**Figure 4.**
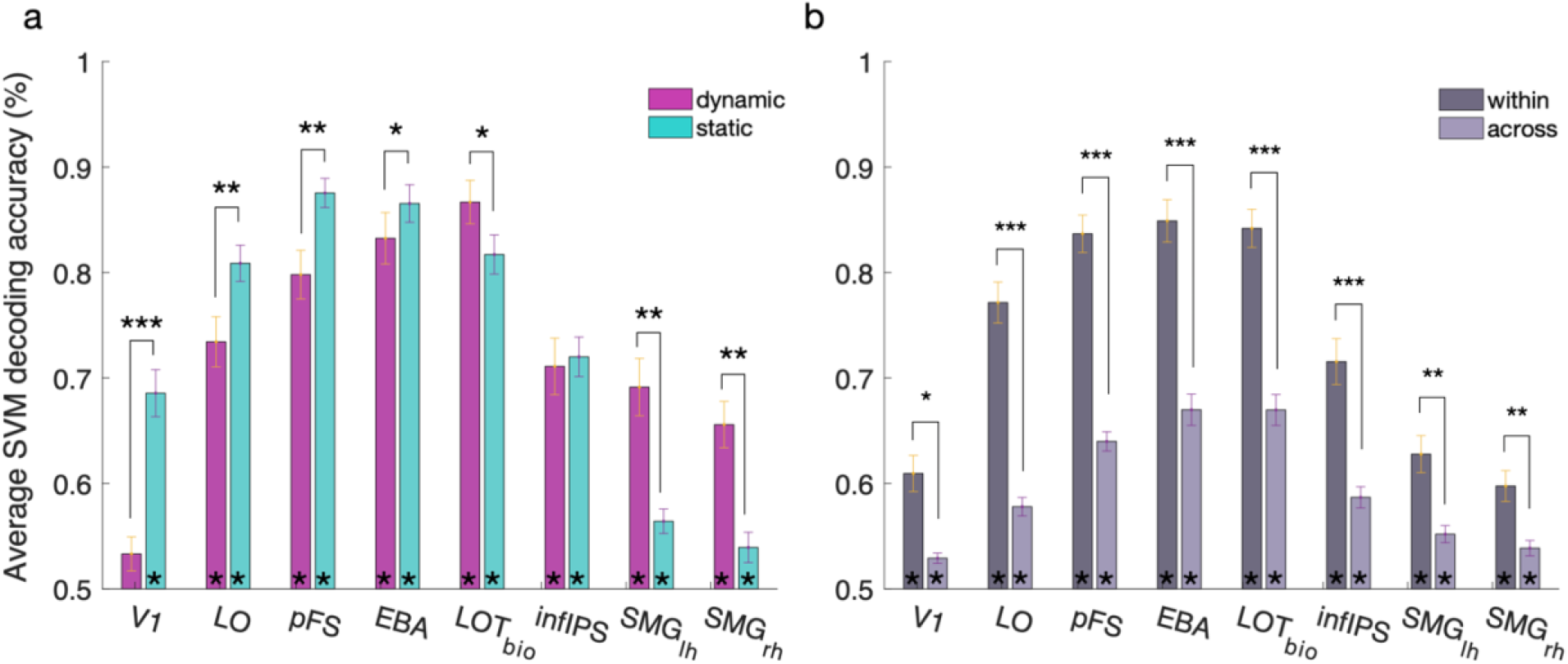
Object category SVM decoding accuracies in each ROI. a) Average SVM decoding accuracies when training and testing within the dynamic (pink) and static (teal) conditions. Asterisks within the bars represent significance in t-tests against chance. All average decoding accuracies were significantly above chance except for the dynamic condition in V1. Asterisks above bars represent paired t-tests across format. In all regions but infIPS, accuracies were significantly higher for one of the formats—LO, pFS, and EBA had significantly higher accuracy in the static condition while LOT-biomotion and bilateral SMG had significantly higher accuracy in the dynamic condition. b) The within stimulus format decoding accuracies, depicted in dark grey bars, were produced by averaging the dynamic and static decoding accuracies in A. The cross-format decoding accuracies are shown in light grey bars. Cross classification was significantly above chance in all regions of interest. Within classification was significantly higher than cross classification in all regions of interest. Error bars represent standard errors. Asterisk notation: * *p* < 0.05, ** *p* < 0.001, *** *p* < 0.0001.

To ensure that the significant decoding of object category from dynamic information was due to differences in the responses to object categories and not contingent upon optic flow information differences that were confounded with category in our stimulus set, we performed a control analysis in which we correlated the dynamic stimulus information with the multivariate fMRI responses (see Methods). No significant positive correlations were observed for any of the regions of interest (*t*s < 2.8, *p*s > 0.06).

We next used a cross-classification method to determine if abstract responses to object categories irrespective of stimulus format exist in our ROIs. The SVM classifier was trained in one stimulus format and then tested in the other format. Decoding accuracies when training on static and testing on dynamic and training on dynamic and testing on static were averaged to produce the light grey bars shown in Figure 4b. We also calculated the within-classification accuracy for training and testing within stimulus format (dark grey bars in Figure 4b; average of the two bars in Figure 4a). Significant cross-classification was observed in all regions of interest (*t*s > 5.31, *p*s < 0.0001, Cohen’s *d*s > 1.37), and was significantly lower than within-classification in all ROIs (*t*s > 5.24, *p*s < 0.0001, Cohen’s *d*s > 1.35). This suggests that the information about object categories in the multivariate pattern responses to the dynamic and static stimuli was sufficiently similar to allow for significant decoding in one stimulus format after being trained on the other.

To further visualize the similarity between the fMRI responses to the object categories in the dynamic and static conditions, we calculated the pairwise Euclidean distances between the patterns of responses to the 6 object categories and the two stimulus formats in each ROI. We then performed a multidimensional scaling analysis on the resultant dissimilarity matrix and visualized the first two dimensions in each of the regions of interest (Figure 5). In all regions, there was a clear separation between the responses to the dynamic (shown in purple and pink) and static stimuli (shown in green and teal). In addition, the ventro-temporal regions and inferior parietal cortex showed a separation amongst the individual object categories. The nearly parallel lines connecting the dynamic and static conditions of the same category indicate that categories with responses that were similar to each other in one condition were also similar to each other in the other condition and is in line with the results of the cross-classification analysis performed earlier. In bilateral supramarginal areas, this object category separation was evident for the dynamic stimulus responses, but the static stimulus responses remained clustered together. In V1, while there was a separation between dynamic and static, the arrangement of categories does not appear to be consistent across conditions.

**Figure 5.**
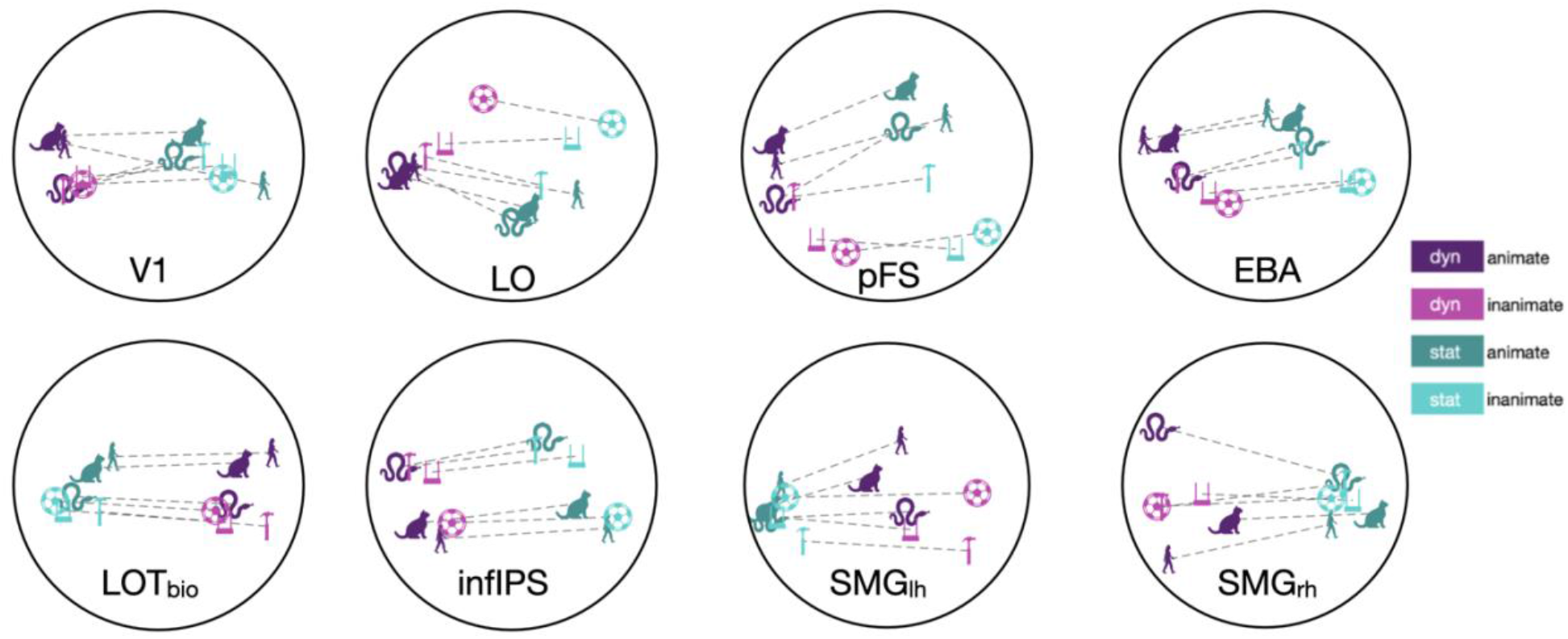
Multidimensional scaling visualization of fMRI response similarity between the object categories presented in the dynamic and static formats. MDS was performed on the similarity matrix obtained from the Euclidean distances of response patterns for the 12 conditions in each ROI. Dotted lines connect dynamic and static presentations of the same object category. The dynamic condition is signified by purple and the static condition is signified by green. Within each condition, the darker hues represent the animate categories while the lighter hues represent the inanimate categories. The 6 object categories are symbolized as with the following icons: human (person from side profile), mammal (cat), reptile (snake), tool (hammer

### Odd-one-out behavioral experiment

To investigate how the responses of each ROI to the 6 object categories in each format relates to the behavioral measure of similarity we performed two behavioral experiments on Amazon Mechanical Turk in which we showed participants three objects (either in static condition or in dynamic condition) and asked them to judge the similarity between the three objects and pick the odd-one-out. We calculated two dissimilarity matrices based on the responses, one for the static stimuli and one for the dynamic stimuli (see Methods). We then averaged the individual object distances from each category to obtain dissimilarity scores between the 6 object categories for the two stimulus formats (Figure 6a). The reliability of these similarity judgments was evaluated for each stimulus format separately (see Methods). Participants rated both stimulus formats with highly stable similarity judgments (*r* = 0.98 for both dynamic and static stimuli). We used multidimensional scaling on the pairwise dissimilarities of each stimulus format to visualize the distance between object categories in the first two dimensions (Figure 6b).

**Figure 6.**
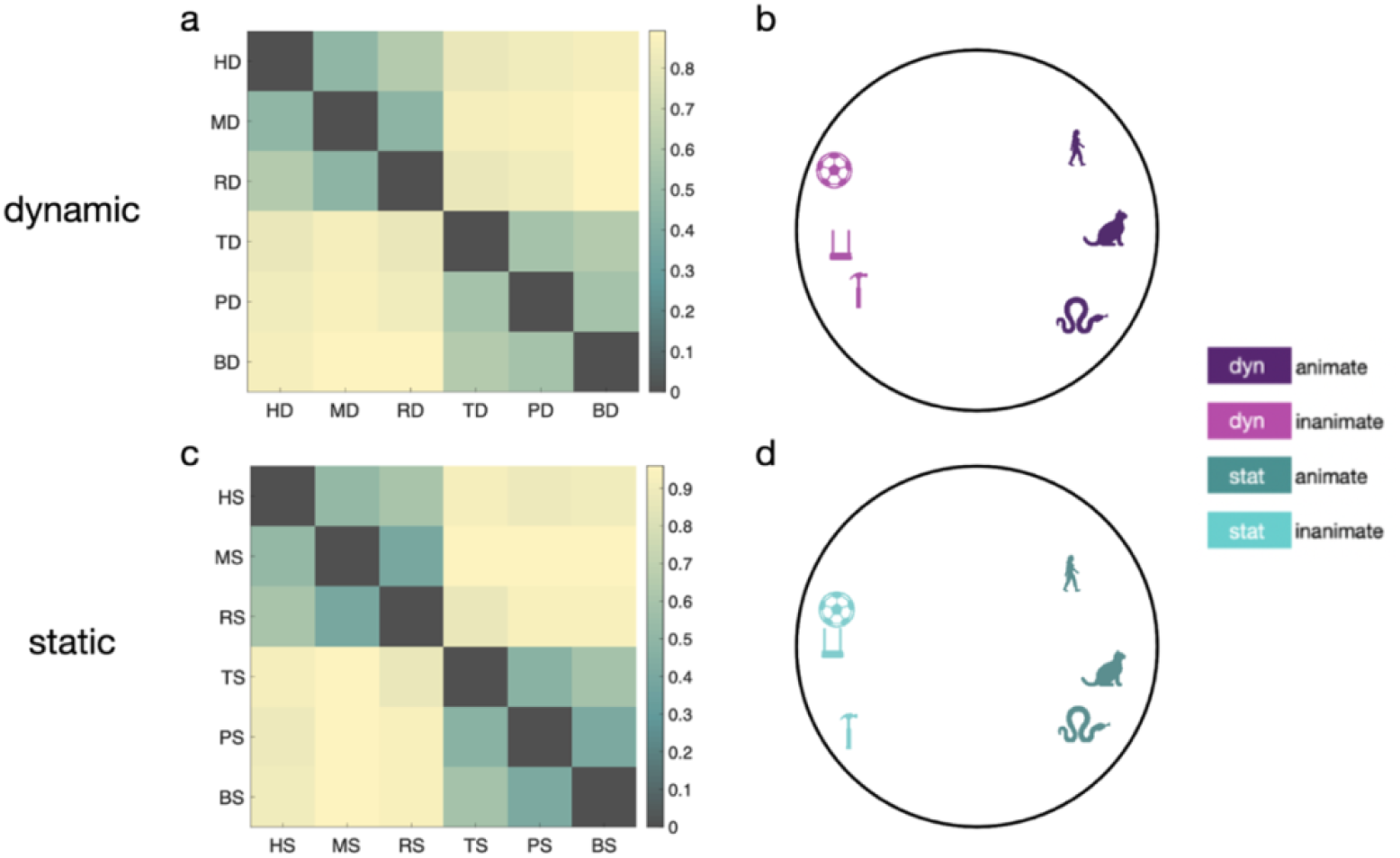
Odd-one-out similarity judgements of dynamic and static stimuli at the category level. The matrices depict pairwise dissimilarity scores between object categories in dynamic (a) and static (c) stimulus formats. The circle plots represent the object categories project into the first two dimensions from multidimensional scaling on their dissimilarities in the dynamic (b) and static (d) stimuli.

The dynamic and static similarity judgments had highly similar structure, showing a clear separation between animate and inanimate categories in the first dimension. The animate (human, mammal, and reptile) and inanimate (tool, pendulum/swing, and ball) categories were also separated from each other along the second dimension in both tasks. Overall, the dissimilarities from the dynamic and static tasks were highly correlated (*r* = 0.98, *p* = 2.80e-10), however, there also appeared to be slight qualitative differences in the arrangement of the inanimate object categories along the second dimension.

To further explore the similarity structure of the dynamic and static stimuli at the exemplar level, a hierarchical clustering algorithm was used on the odd-one-out similarity judgments (Figure 7). Similar to the MDS of odd-one-out judgements at the category level, a gross distinction between animate and inanimate objects was observed for both the static and dynamic conditions. Moreover, as in the MDS, the three object categories within the animate and inanimate superordinate categories are largely distinguished in both formats. However, the clustering algorithm also revealed several interesting differences in the similarity judgments of the same objects when presented in either static image or dynamic optic flow format. For example, the dynamic baboon stimulus, a clip of a baboon sitting and feeding, was grouped with the human stimuli, while the static baboon stimulus was grouped with the mammal stimuli. Similarly, the dynamic presentation of the two pendulum stimuli were grouped with the swings, presumably due to their shared movement patterns, while their static presentations were grouped with the balls, likely due to their shared global form. These deviations of specific exemplars from their category clusters illustrate important differences in the category information provided by dynamic and static visual cues and shed light on some of the heuristics that are used to guide similarity judgments in the absence of either form or motion information. When luminance-defined edges are not available, robust category information can be derived from dynamic motion-isolated inputs.

**Figure 7.**
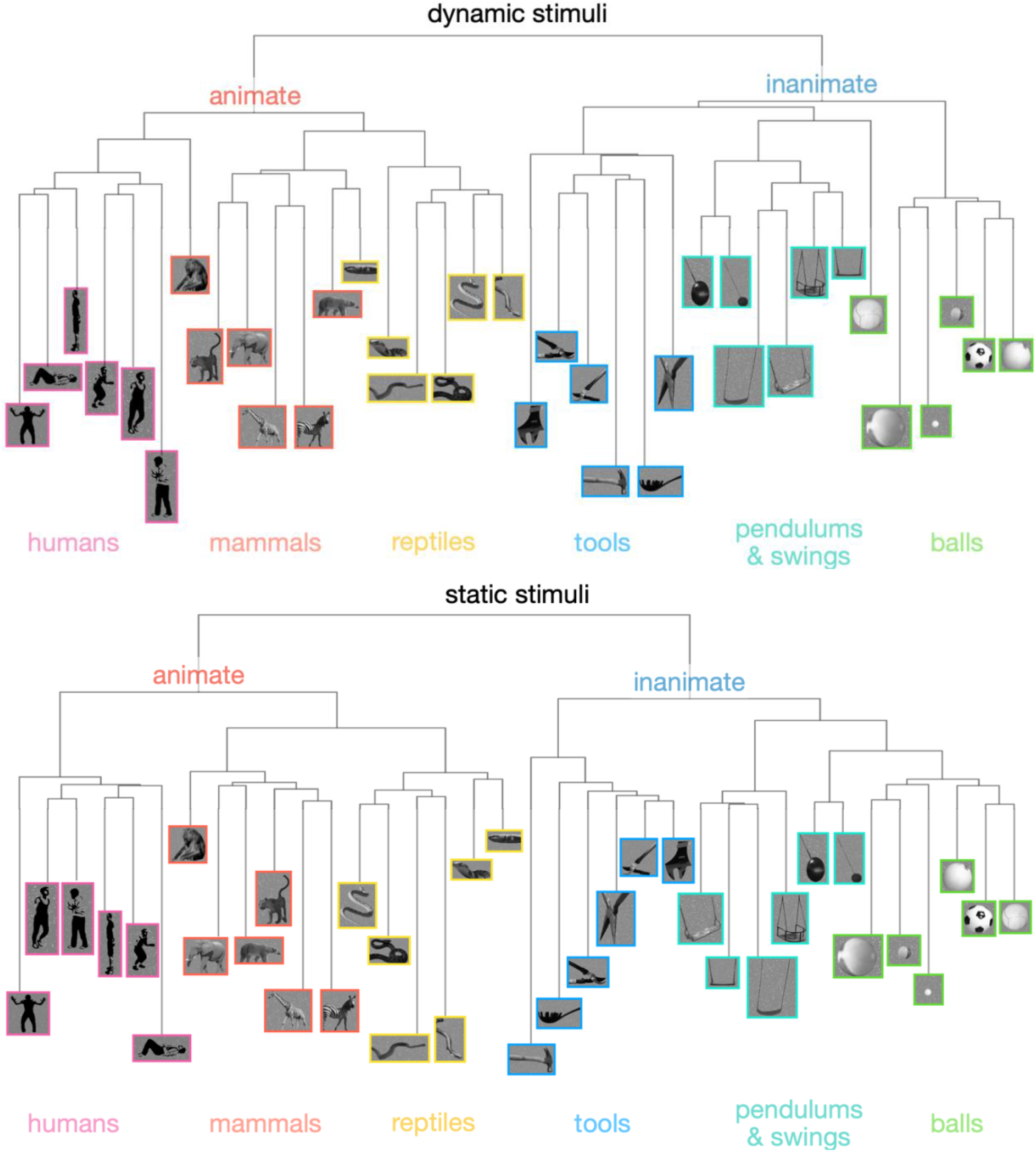
Hierarchical clustering of odd-one-out similarity judgments of the dynamic and static stimuli at the exemplar level. Edited versions of the static stimuli were used to visualize the similarity structure of both the dynamic (top) and static (bottom) stimuli as category of the dynamic stimuli cannot be gleaned from individual frames. The scale and position of the objects are not representative of the stimuli during presentation. Stimulus borders were colored to distinguish the six object categories. The human stimulus examples were modified into two-tone images for this figure to deidentify the individuals in the stimuli.

To investigate how the object category fMRI responses to each format relate to behavioral judgements of similarity, we correlated the dissimilarity scores from the dynamic and static behavioral experiments (dynamic and static reliability: 0.985) to those obtained from the Euclidean distances between the multivariate response patterns in each region of interest (*r*s: dynamic > 0.03; static > 0.02, apart from right SMG, see below). As shown in Figure 8, most ventral and lateral temporal regions—LO, pFS, EBA, LOT-biomotion—showed significant correlations with the object similarity judgments for both the dynamic and static stimuli (dynamic: *p*s < 0.01; static: *p*s < 0.05). The responses in infIPS were not correlated to object similarity judgments for either the dynamic or static stimuli (dynamic: *p* = 0.12, static: *p* = 0.59). The activity in left SMG was significantly correlated with the similarity judgments for the dynamic stimuli (*p* = 0.001), but not for the static stimuli (*p* = 0.59). Similarly, the activity in V1 was significantly correlated with similarity judgments for the static stimuli (*p* = 0.02), but not for the dynamic stimuli (*p* = 0.14). The only significant difference between the correlations of the behavioral similarity judgments and the fMRI responses to the two conditions was found in the left SMG area, in which the correlation was significantly higher with similarity judgments of the dynamic stimuli compared to the static stimuli (*t*(14) = 3.32, *p* = 0.04, Cohen’s *d* = 0.86). In the right SMG area, the *r* value was -0.0083 for the static condition, signifying a reliability of zero. As this suggests that the responses to the static stimuli in this region were unreliable, the correlation between the multivariate fMRI responses in the right SMG to the static stimuli with behavioral assessments of their similarity will not be interpreted.

**Figure 8.**
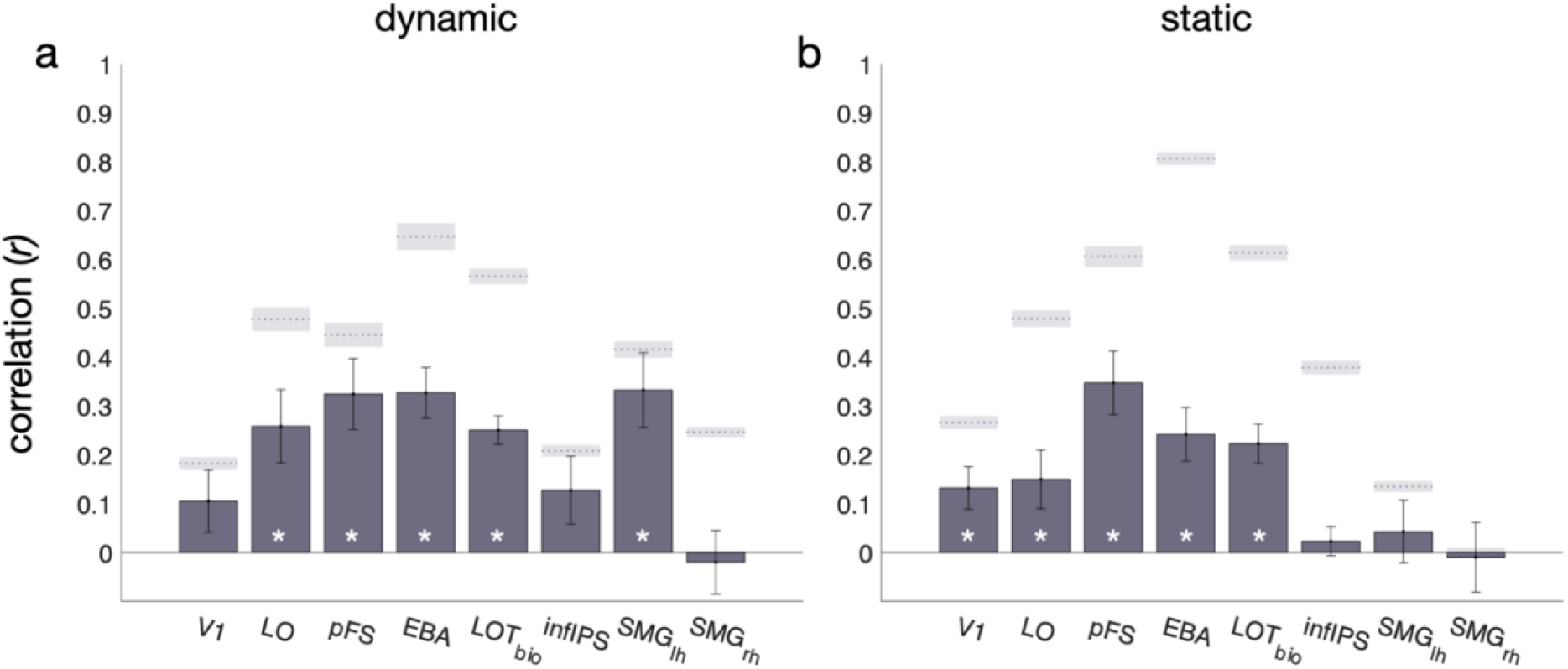
Correlation of Euclidean distance between multivariate fMRI responses and behavioral dissimilarity matrices for a) dynamic and b) static stimuli. * *p*s < 0.05. Error bars represent standard errors. Shaded regions represent the average noise ceiling (dotted line) and the standard error of noise (shaded region) for each ROI.

## Discussion

Motion is an important visual cue that can provide category-relevant information in the absence of luminance-defined edges and form. Here, we introduce a novel approach to systematically separate form and motion signals and study the contribution of the motion signal to object category processing in isolation. To our knowledge, our study is the first to use this approach to compare the neural processing of form and motion signals from several animate and inanimate object categories. We sought to determine whether category-relevant information from the two sources is shared across the visual system by comparing dynamic and static category processing in regions of interest across visual occipito-temporal and parietal cortices. The two highly dissimilar information sources produced distinct but overlapping representations of animate and inanimate object categories, with a shift in processing primarily static information in more ventral regions to primarily dynamic information in more dorsal regions of cortex.

### Categorizing Objects with Motion Information

An object identification task was used to determine whether our method for simulating the extracted motion information in dynamic flow fields could produce stimuli in which objects were recognizable. Our findings illustrate that, not only do people categorize motion-defined *animate* objects with high accuracy (Pinto, 2006; Pinto, 1994; Pavlova et al., 2001), this high performance also holds for three *inanimate* object categories: tools, swinging objects, and balls. These results extend previous research by showing that a wide range of objects spanning animate and inanimate categories can be recognized from just motion information. Our odd-one-out judgment task further demonstrated that the similarity judgments for the dynamic and static stimuli were highly correlated. This consistency suggests that people infer the similarity of objects from the two sources of information in a similar way.

When discussing the perception of objects from motion, it is important to distinguish between two types of information that can be gleaned from motion cues: 1) structure from motion, a percept of a form arising from the global integration of coherent local motion vectors, and 2) types of actions that are diagnostic of a particular object category such as walking, swinging, tool use, bouncing, etc. Though it was not within the scope of this study to systematically distinguish these two sources, the exemplar level clustering of our odd-one-out data qualitatively suggests that both factors may play an important role in subjects’ judgements of object similarity. For example, images of pendulums and bouncing balls maybe judged to be similar since they both contain a round shape, but distinct in dynamic form because they move differently.

### Format-dependent processing of object categories

Comparison of the object category information across the two stimulus formats revealed differences in many of our regions of interest. Our findings suggest that stimulus format matters for: 1) processing of animate and inanimate objects—indicated by the regions of interest with significant interactions between stimulus format and univariate animacy preference (i.e., pFS and left SMG)—and 2) discriminating object categories within format—indicated by regions with significant differences in the multivariate classification accuracy of the responses to dynamic and static stimuli (i.e., all regions but infIPS). Broadly speaking, we found that the most ventral and posterior regions we examined (LO, EBA, and pFS) showed higher classification in the static condition, while most dorsal and anterior regions (LOT-biomotion and bilateral SMG) had stronger classification in the dynamic condition. Interestingly, infIPS used both sources of information without dominance of one source over the other. Importantly, all regions of interest but V1 showed robust responses to, and significant decoding accuracies of, all categories presented in both static image and dynamic motion formats. Thus, differential multivariate processing of object category based on stimulus format in these regions is a matter of degree. These results align with predictions from the model presented by Giese and Poggio (2003), in which form and motion signals are processed by distinct neural populations that largely overlap in topographic regions across ventral and dorsal cortex.

### Animate and Inanimate Category Processing

Relative to static images, investigation of topographic organization of object category processing driven by motion information has been largely neglected. However, an important exception can be found in the work of Beauchamp and colleagues (2003), in which they compared univariate fMRI responses between 1) full form videos and static images of humans and tools and 2) full form videos and point-light displays of humans and tools. Beauchamp et al. (2003) argued for two processing pathways—form and motion. Lateral temporal regions (STS and MTG), respond to their preferred category, humans and tools, respectively, in both PLDs and videos, suggesting category preference from motion without requiring form. Meanwhile, ventral temporal cortex (lateral and medial fusiform), needed form information for category preference responses. Our results are in agreement with these findings and demonstrate that the topography of animacy preference is not dependent on or exclusive to the human and tool categories—it also expands to other animate objects such as mammals and reptiles, and other inanimate objects such as pendulums/swings, and balls. These results suggest that large-scale animacy preference maps (Konkle & Caramazza, 2013, Sha et al., 2015) found with static objects in the brain might also be present for motion defined stimuli. Future studies with a larger stimulus set and sufficient power to perform whole-brain analyses will be crucial for expanding our findings beyond functionally defined regions of interest in VOTC and parietal cortex.

### Distinct but Overlapping Representations of Object Category for Dynamic and Static Stimuli

Using linear SVM classifiers, we decoded object category with high accuracy in all regions tested. In all regions but V1 and the right supramarginal area, both information sources drove object representations that were sufficiently distinguishable from each other to allow for high classification performance. Extracting form and motion information from the same objects and presenting them separately also allowed us to investigate the extent to which the representations are overlapping across stimulus formats. We used a cross-classification approach to identify regions that have format independent responses. A similar analysis has been used previously to study fMRI responses to human actions in full form videos and images (Hafri et al., 2017). Our results are largely in qualitative agreement with those of Hafri and colleagues, with the exception that we found significantly more widespread cross-classification, possibly because our static stimuli were source matched to our dynamic stimuli. Cross-decoding in all regions (apart from V1) suggests that the object category representations driven by static and dynamic information were sufficiently distinct to allow for significant within format classification, but also sufficiently overlapping that their shared information could lead to significant cross-classification. These results suggest the existence of abstract object category responses that pool information about object category across various cues in the visual input.

### Relationship between brain and behavior

Multivariate responses to both the dynamic and static conditions in LO, pFS, EBA, and LOT-biomotion—the ventral and lateral regions—were correlated with the object similarity judgments of the dynamic and static stimuli, respectively, with no differences across condition. This implies that the fMRI responses in these regions follow the structure of the stimulus similarity characterized by our odd-one-out experiment. The only region to show a difference in correlation across the stimulus conditions was the left supramarginal area, which showed higher correlations for the fMRI responses to the dynamic relative to the static stimuli. By contrast, the right supramarginal area showed no significant correlation to behavioral judgments of either condition, which indicates a lateralization of inanimate category processing to the left supramarginal area. This left lateralization has been shown previously in research on tool processing (Beauchamp et al., 2003). Importantly, not all regions that showed significant animacy preference or object category decoding had responses that were significantly correlated with the similarity structure of the behavioral judgments. In V1 and infIPS, the fMRI responses to both conditions were unrelated to the similarity judgments of both stimulus types, suggesting that these regions were extracting features irrelevant to similarity judgments on the objects.

### Conclusion

In sum, our study demonstrates that in regions across occipito-temporal and parietal cortices, category responses driven by isolated motion signals parallel category responses to static form signals in a number of interesting ways. Regions that are traditionally considered part of the visual object recognition pathway that processes static information, such as the pFS, LO, and EBA, also extract robust object category information from isolated motion signals relevant to behavioral judgments of object similarity. Furthermore, cross-classification of object categories in all regions suggests that object-category information from static and dynamic signals overlap. Lastly, preferential processing of certain kinds of objects, such as animate or inanimate objects, is sensitive in some regions, i.e., the pFS and left SMG, to the format of visual information. Using the stimulus generation approach we have introduced, future studies can expand beyond the six object categories tested here and introduce parametric manipulations of dimensions that are likely to play an important role in differential processing of motion-derived object categories. Candidate dimensions include the type of action or movements that the objects are performing as well as the orientation from which the movements are viewed. Such studies will be important for furthering our understanding of how various visual cues to object-category are processed and integrated together to form rich and robust object representations in the human brain.

## Acknowledgments

We thank the late and great Leslie G. Ungerleider for her mentorship and guidance throughout this project, Chris Baker for insightful feedback, and Julian De Freitas for inspiring discussions that helped in forming the initial interest in this research area. This research was supported by the National Institute of Mental Health Intramural Research Program (ZIA-MH-002909).

## Notes

**Conflict of interest statement:** The authors declare no competing financial interests.

### Competing Interest Statement

The authors have declared no competing interest.

### Summary of Updates

Figure 7 fixed

